# Genetic demultiplexing and transcript start site identification from nanopore sequencing of 10x Genomics multiome libraries

**DOI:** 10.64898/2026.03.31.715454

**Authors:** J. Mears, P. Orchard, A. Varshney, M. Bose, C.C. Robertson, M. Piper, E. Pashos, V. Dolgachev, N. Manickam, P. Jean, D.W. Kitzman, E.B. Fauman, F. Damilano, R.J. Roth Flach, B. Nicklas, S.C.J. Parker

## Abstract

Short-read Illumina sequencing of 10x Genomics single-nucleus multiome libraries captures only the 3’ end of RNA transcripts, losing transcription start site (TSS) information. Here we demonstrate nanopore sequencing of 10x multiome libraries, which enables the profiling of full length transcripts. We show concordance with common short-read sequencing based workflows including successful genetic demultiplexing of nanopore data despite its higher error rate. We compare TSS identified using nanopore sequencing of multiome cDNA to those identified using a short-read 5’ assay, and provide an optimized approach for the preprocessing of nanopore reads prior to TSS identification. We find that nanopore sequencing of multiome cDNA captures a median of 63% of the TSS detected by the 5’ assay.

## Background

Single-cell and single-nucleus experiments have grown tremendously popular in the last decade. 10x Genomics is one commonly-used platform, providing assays for different types of single nucleus multi-omics profiling including 3’ gene expression (3’ GEX), 5’ gene expression (5’ GEX), and multiome (3’ GEX and chromatin accessibility (ATAC)). Profiling of 3’ or 5’ gene expression involves the generation of barcoded, full transcript length cDNA libraries, which are fragmented prior to short-read sequencing, resulting in sequencing reads representing either the 3’ or 5’ end of a transcript.

Thanks to decreasing sequencing error rates and increasing throughput^1–3^, nanopore-based long-read sequencing technologies such as the Oxford Nanopore Technologies (ONT) platform have become a viable option for sequencing 10x libraries^4–8^. Long-read sequencing of the full-length (unfragmented) cDNA produced during 10x library preparation enables recovery of information lost in the standard short-read workflow; for example, 5’ transcript ends can be inferred from the 3’ gene expression library cDNA, enabling detection and quantification of transcription start sites (TSS) without the need for separate profiling with a 5’ GEX assay.

However, the application of long-read ONT sequencing to single cell assays is still relatively new. To date, most studies have focused on the use of ONT sequencing to identify isoform expression patterns^5,9–22^. While many technical aspects of analysis of ONT sequencing have been well characterized such as barcode/UMI identification and correction^23–25^, other challenges remain. We enumerate two such examples here. First, many short-read single cell studies leverage genetic multiplexing -- pooling samples from different donors prior to loading onto the 10x platform -- to improve doublet detection, enable higher loading concentrations, and decrease cross-sample technical variance^26–30^. This approach relies upon the ability to genetically demultiplex in downstream analysis (assigning nuclei to donors on the basis of genetic variation observable in sequencing reads). Therefore, it is unclear whether this technique will be as reliable in a study design using ONT sequencing, which has a higher sequencing error rate than Illumina short-read sequencing. Second, technical differences in transcript capture and transcript completeness have been previously reported between 10x Genomics 3’ and 5’ gene expression assays^4,6^, raising the question as to how similar TSS detected using ONT sequencing of multiome gene expression will be to those detected using short-read sequencing on a 5’ GEX assay.

Here we demonstrate long-read (ONT) sequencing of 10x multiome RNA modality libraries shows high concordance with common short-read (Illumina) sequencing-based workflows including successful genetic demultiplexing of ONT sequencing data. We additionally compare the TSS identified using ONT sequencing of multiome libraries to those identified using short-read sequencing of 5’ GEX libraries, and outline an optimized approach for the preprocessing of ONT reads and removal of artifact signals prior to identification of 5’ transcript ends from 3’ GEX cDNA.

## Results and Discussion

We isolated nuclei from 60 frozen vastus lateralis skeletal muscle tissue biopsies (52 unique donors), processing 20 samples per batch, resulting in three pools of nuclei that each comprised 20 samples (20 unique donors) (Fig. 1A, left). From each pool we generated two single nucleus multiome libraries, resulting in six total single nucleus multiome libraries (Fig. 1A, middle). Both multiome modalities (3’ GEX and ATAC) underwent short-read sequencing, and the intermediate full-length cDNA from the 3’ GEX library preparations underwent long-read ONT sequencing (ONT GEX) (Fig. 1A, right). Additional nuclei from the same three nuclei pools were used to generate a total of six 5’ GEX libraries, which underwent short-read sequencing (Fig. 1A, middle and right). Pseudobulk gene expression showed high correlation between ONT GEX and 3’ GEX (Fig. 1B).

**Fig. 1.**
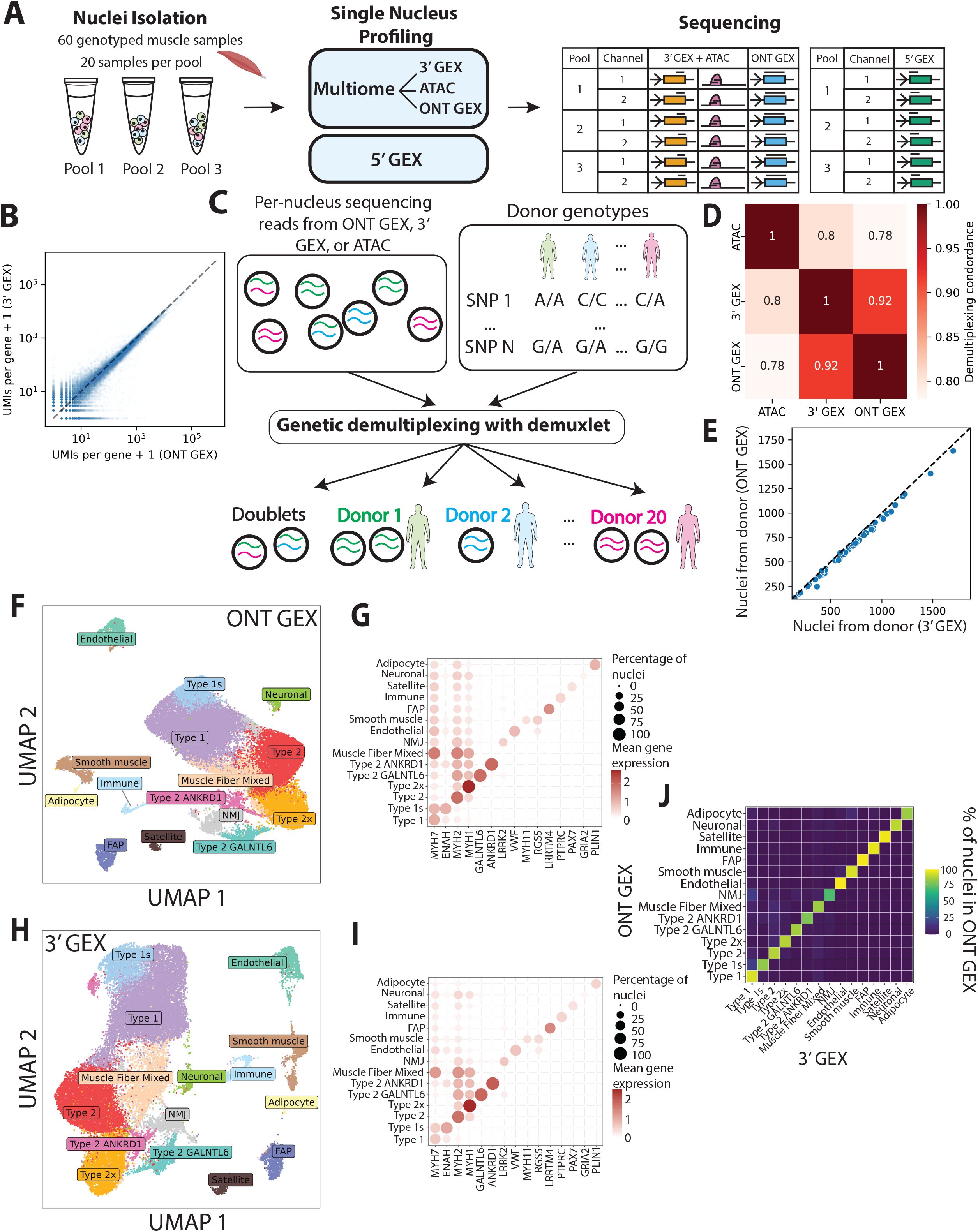
Experimental overview and comparison to 3’ GEX. **(A)** Overview of experimental design. **(B)** Pseudobulk gene expression in ONT vs Illumina-sequenced 3’ GEX libraries. Each point represents one gene. **(C)** Overview of genetic demultiplexing process. We used demuxlet to perform genetic demultiplexing using sample genotypes and sequencing reads from either ONT GEX, 3’ GEX, or ATAC reads. **(D)** Concordance in barcode demuxlet assignments between ONT GEX, 3’ GEX, and ATAC reads. For a given barcode, the assignment is considered concordant if the barcode is either marked as a doublet in both datasets, or marked as a singlet with the same inferred donor in both datasets. **(E)** Number of nuclei assigned to each donor when demultiplexing with ONT GEX vs 3’ GEX reads. **(F)** Clustering of ONT GEX data and projection in UMAP space with nuclei cell type assignments. **(G)** Mean ONT GEX expression and percentage of nuclei expressing key markers across nuclei types. **(H)** Clustering of 3’ GEX data and projection in UMAP space with nuclei cell type assignments. **(I)** Mean 3’ GEX expression and percentage of nuclei expressing key markers across nuclei types. **(J)** Concordance in cell type assignments between ONT GEX and 3’ GEX. Heatmap values are expressed as the percentage of nuclei from the ONT GEX data with a matching cell type label in the 3’ GEX data.

Genetic demultiplexing, which entails assigning nuclei to donors on the basis of genetic variants observed in sequencing reads, is commonly employed in single nucleus experiments (Fig. 1C). Because it relies upon accurate detection of genetic variants in sequencing reads, demultiplexing is expected to become less reliable as sequencing error rate increases. As ONT sequencing has higher error rates than Illumina sequencing, we asked whether demultiplexing was still feasible using the ONT GEX data. To investigate this, we used demuxlet^26^ to perform genetic demultiplexing using ONT GEX, 3’ GEX, or ATAC reads, and compared demuxlet’s donor assignments and doublet calls across the datasets. Overall, concordance was highest between 3’ GEX and ONT GEX (92%; Fig. 1D). As expected, the number of nuclei identified per donor correlated strongly across the sequencing platforms, though the number of nuclei per donor was slightly lower for ONT GEX than 3’ GEX (Fig. 1E). These results suggest that ONT-sequenced snRNA-seq libraries can be successfully demultiplexed, with perhaps a slight decrease in the number of nuclei identified per donor relative to comparable short-read data.

After QC and demultiplexing, we recovered 32,468 high-quality nuclei across the six multiome libraries, and 61,565 nuclei across the six 5’ GEX libraries (Fig. S1A). The number of UMIs per nucleus varied across individual libraries but was not systematically different across the modalities (Fig. S1B, S1C). We clustered the ONT GEX and 3’ GEX and found they yielded highly similar clustering results (Fig. 1F-J), confirming that ONT sequencing of 3’ GEX cDNA from a multiome experiment recapitulates key gene-level information.

One advantage of long-read sequencing of multiome libraries is the ability to identify transcript 5’ ends, enabling detection and quantification of TSS. However, as differences in transcript capture and completeness have been previously reported between the 10x Genomics 3’ and 5’ gene expression assays, TSS identified using ONT GEX may differ somewhat from those identified using short-read sequencing with 5’ GEX. Therefore, we compared the two. One challenge in TSS identification is that some apparent 5’ ends may be artifacts, for example due to template-switch oligo (TSO)-related strand invasion as described elsewhere^31^. A previously-published tool to identify and quantify 5’ ends while adjusting for these artifacts is the Single Cell Analysis of Five-prime Ends (SCAFE) software package^32^. SCAFE accepts a BAM file, extracts inferred transcript 5’ ends, removes likely artifactual 5’ ends, clusters 5’ ends into prospective elements called ‘transcribed cis regulatory elements’ (tCREs, which are a proxy for TSS), and filters the prospective tCREs to a high-quality list based on indicators such as chromatin accessibility over tCREs and the presence of unencoded guanines on transcript ends indicative of 5’ capping (Fig. 2A, middle). tCREs are expected to be highly enriched near active gene TSS. We ran SCAFE on ONT GEX and 5’ GEX datasets and assessed the quality of the resulting tCRE calls by examining tCRE overlap with previously-published skeletal muscle chromatin states^33^. To avoid confounding due to the differing number of nuclei in the ONT GEX and 5’ GEX libraries, we subsampled matched ONT GEX - 5’ GEX pairs to the same number of nuclei. SCAFE encounters a read length-related error when run on single nucleus long-read data; upon fixing this in the naive manner (trimming the 3’ ends of aligned reads to reduce the read length), we found that most tCREs discovered using ONT GEX overlapped tCREs discovered using 5’ GEX (Fig. 2B, S2, S3), and tCREs from both ONT GEX and 5’ GEX overlapped active TSS and flanking active TSS chromatin state regions, as expected (Fig. 2C, S4, S5). However, for some libraries, the trimmed ONT GEX tCRE calls included a substantial number of tCREs in quiescent regions and in the weak and strong transcription chromatin states (which generally occur over gene bodies and should not contain many transcribed regulatory elements; Fig 2C, 2D, S4, S5), likely representing false positive tCREs. In such cases we found additional filtering of the ONT read alignments prior to running SCAFE (Fig. 2A, left) substantially reduced the number of tCRE calls in quiescent regions and transcribed regions with only a slight reduction of tCRE calls near active TSS (Fig. 2C, S4, S5). In brief, this filtering removes alignments missing a short expected residual TSO sequence at the 5’ end or having excessive softclipping at the 5’ end, and identifies and filters cases in which apparent PCR duplicates nominate discrepant 5’ end positions. We provide scripts implementing this filtering for use with ONT BAM files from the official ONT pipeline prior to running SCAFE (https://github.com/porchard/pre-SCAFE-processing).

**Fig. 2.**
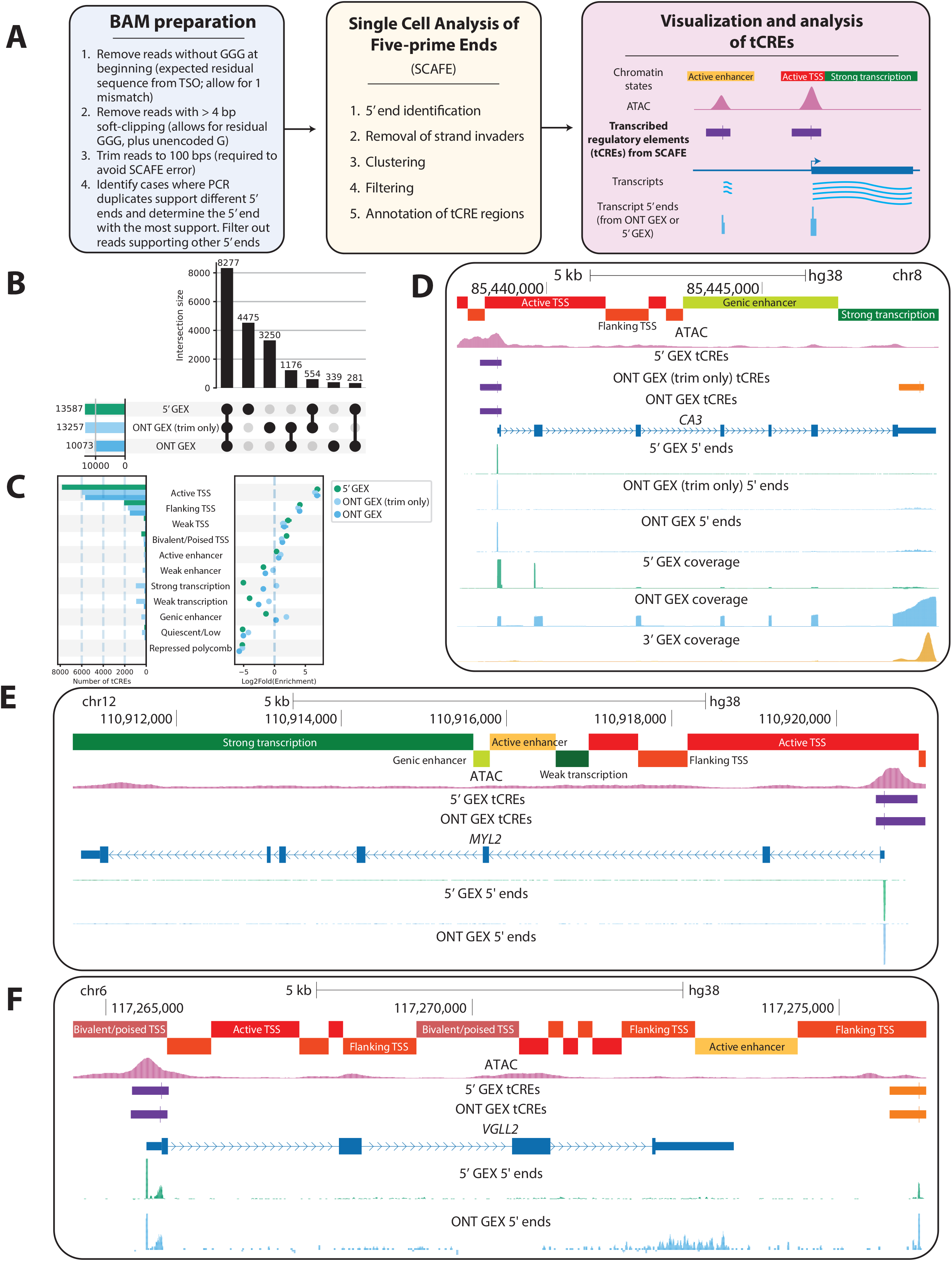
tCRE discovery and comparison to 5’ GEX. **(A)** Left panel: Overview of processing steps applied to ONT alignments before running SCAFE. Middle panel: Overview of key steps in the SCAFE method. Right panel: SCAFE output contains annotated bed files of transcribed cis-regulatory elements (tCREs) at the 5’ ends of sequenced RNA molecules. **(B)** Overlap between tCREs obtained from SCAFE using 1) ONT GEX with only basic trimming rather than the full preprocessing outlined in the left panel of Fig 2A (“ONT GEX (trim only)”), 2) ONT GEX with the full pre-processing described in the left panel of Fig 2A (“ONT GEX”), or 3) 5’ GEX. **(C)** ChromHMM annotated chromatin state overlap (left) and enrichment (right) with sets of tCREs obtained from SCAFE using 1) ONT GEX with only basic trimming rather than the full preprocessing outlined in the left panel of Fig 2A (“ONT GEX (trim only)”), 2) ONT GEX with the full pre-processing described in the left panel of Fig 2A (“ONT GEX”), or 3) 5’ GEX. **(D)** UCSC Genome Browser, showing ChromHMM chromatin state annotations, ATAC signal, tCREs from 5’ GEX / ONT GEX (trim only) / ONT GEX, 5’ ends from 5’ GEX / ONT GEX (trim only) / ONT GEX, and sequencing coverage in 5’ GEX / ONT GEX / 3’ GEX at the *CA3* locus. SCAFE identifies a tCRE at the *CA3* gene TSS using 5’ GEX / ONT GEX (trim only) / ONT GEX ONT, and a tCRE in a strong transcription region at the gene 3’ end using ONT GEX (trim only). **(E)** UCSC Genome Browser, showing ChromHMM chromatin state annotations, ATAC signal, tCREs from 5’ GEX / ONT GEX, and 5’ ends from 5’ GEX / ONT GEX at the *MYL2* locus. SCAFE identifies a tCRE using ONT GEX and 5’ GEX corresponding to the *MYL2* promoter region. **(F)** UCSC Genome Browser, showing ChromHMM chromatin state annotations, ATAC signal, tCREs from 5’ GEX / ONT GEX, and 5’ ends from 5’ GEX / ONT GEX at the *VGLL2* locus. SCAFE identifies a tCRE using ONT GEX and 5’ GEX corresponding to the *VGLL2* promoter region, as well as a tCRE downstream of the gene in both modalities.

Comparing the final set of tCREs obtained from our ONT GEX data to tCREs obtained using the 5’ GEX assay, we found that most tCREs captured using the 5’ GEX assay are also captured by ONT GEX (Fig. S2, S3). These similarities are evident when examining individual loci such as the promoter of the gene *MYL2* (Fig. 2E), which encodes an important protein for muscle contraction and is highly expressed in skeletal muscle, or the *VGLL2* locus (Fig. 2F). Nevertheless, ONT GEX fails to detect a considerable number of tCREs detected by 5’ GEX in active TSS and flanking active TSS chromatin states (Fig. S4). This may reflect differences in the assay kits, imperfections in the ONT library preparation, or the need for further optimization of computational processing, and is in agreement with previous work^4,6^ suggesting differences in transcript capture between 3’ and 5’ GEX assays.

## Conclusions

ONT sequencing of multiome experiments provides an opportunity to assay full length cDNA molecules and recover information beyond that captured using short-read sequencing, including 5’ transcript start information. We demonstrate that ONT sequencing of 3’ GEX libraries successfully recapitulates standard short-read 3’ GEX results in a common study design, including enabling successful genetic demultiplexing. We find it enables detection of many of the same TSS captured using the 10x Genomics 5’ assay, though it is notable that a substantial number of TSS identified with the 5’ assay were not detected in the ONT data. We provide scripts for the preprocessing of 3’ GEX ONT alignments prior to use with the SCAFE tCRE detection tool. As long-read sequencing and single nucleus experimental designs each continue to grow in popularity, we predict these results will be useful in designing and analyzing experiments.

## Methods

### Sample procurement

Samples were frozen needle biopsy samples taken from the vastus lateralis muscle of participants of the UPBEAT study, and the SECRET I^34^ and SECRET II^35^ randomized, controlled clinical trials.

### Nuclei isolation

Skeletal muscle was cryopulverized in a tissue tube using a Covaris CP02 cryoPREP Automated Dry Pulverizer on dry ice. 4-5 tubes of pulverized samples (∼220mg tissue total) were resuspended in 1 mL of lysis buffer (50mM HEPES pH7.5, 140mM NaCl, 1mM EDTA pH 8.0, 10% glycerol, 0.5% NP-40, 0.25% Triton X-100, 5ug/mL 7-AAD, 1U/uL RNase Inhibitor (Sigma Protector RNase Inhibitor)) and pooled together (∼4ml of buffer) in a 7.5ml dounce homogenizer (Kimble Kontes 7ml tenbroek glass homogenizer, Fisher Scientific Fisher: K885000-0007) on ice. Samples were homogenized for 15 strokes and the homogenate was transferred to two 2 mL tubes. A wide bore tip pipette was used to lyse the cells on ice with 9 strokes/min for 5 minutes. Samples were centrifuged at 500 xg for 5 minutes at 4oC. Supernatant was aspirated and pellets were resuspended in 1 mL of ice cold wash buffer (10mM Tris-HCl pH 7.4, 10mM NaCl, 3mM MgCl2, 1mM DTT, 5ug/mL 7-AAD, 1% BSA, 0.1% Tween20, 1U/uL RNase Inhibitor (Sigma Protector RNase Inhibitor)) and centrifuged at 100 xg for 1 minute at 4oC. Supernatant was collected and loose debris pellets were discarded. Supernatant was filtered through 2 sets each of 70 μm, 50 μm, and 30 μm filters, washing with an additional 200 μL of wash buffer after each pass. Filtrate collected in a 15ml tube was centrifuged at 500 xg for 5 minutes at 4oC. After centrifugation the supernatant was removed and the pellets were resuspended in 3000 μL of stain buffer (1%BBSA in PBS+0.1%Tween, and 5ug/mL 7-AAD). Samples were incubated shaking on ice for 15 minutes before filtering with a 20 μm filter. Nuclei were sorted with the MACS quant Tyto (Miltenyi Biotech, Auburn CA). After sorting, 2 μL of nuclei was taken and added to 8 μL of 1%BSA+PBS for nuclei counting and grading. Remaining nuclei were sent to the Advanced Genomics Core at the University of Michigan for library prep and sequencing. Nuclei were isolated according to the above protocol across three batches and each multiplexed library contained nuclei isolated from ∼ 20 donors such that multiple libraries contain nuclei from the same individual.

### Library prep

Multiome (snRNA + snATAC) libraries were prepared using the Chromium Next GEM Single Cell Multiome Reagent Kit A (10x Genomics, PN-1000284) according to the manufacturer’s instructions. (Chromium Next GEM Single Cell Multiome ATAC + Gene Expression, Document Number CB000338 Rev G, 10x Genomics, (2024, October 4).)

For 5’ GEX libraries, single cell suspensions were subjected to counting and viability checks on the LUNA Fx7 Automated Cell Counter (Logos Biosystems) and diluted to a concentration of 700 -1000 cells/ul. Single cell libraries were generated using the 10x Genomics Chromium Controller with 5’ HT Gene Expression v2 reagents following the manufacturer’s protocol (10x Genomics). Final library quality was assessed using the LabChip GX (PerkinElmer).

### Illumina sequencing

The multiome snRNA modality (3’ GEX) was subjected to 28×151bp of sequencing, while the multiome snATAC modality was subjected to 51×51bp paired-end sequencing, according to the manufacturer’s protocol (Illumina NovaSeqXPlus). BCL Convert Conversion Software v4.0 (Illumina) was used to generate de-multiplexed fastq files 5’ GEX was subjected to 151bp paired-end sequencing according to the manufacturer’s protocol (Illumina NovaSeqXPlus). BCL Convert Conversion Software v4.0 (Illumina) was used to generate de-multiplexed Fastq files.

### Nanopore sequencing

The cDNA from the snRNA protocol was cleaned using 1.8X Ampure XP beads prior to ONT library prep. The samples were prepared using the Ligation Sequencing Kit V14 (SQK-LSK114) along with the PCR Expansion Kit (EXP-PCA001). ONT libraries were sequenced on an R10.4.1 PromethION flow cell. Each library was loaded in two attempts, with a nuclease wash performed between runs. Libraries ONT GEX 1 and ONT GEX 2 were basecalled with Dorado version 7.2.13 (high accuracy model), while all other libraries were basecalled with Dorado 7.3.11 (fast model).

### Multiome data processing

The gene expression modality of the single nucleus multiome data was processed using STARsolo^36^ (v. 2.7.10a, with hg38 fasta file and GENCODE v30-derived genome annotations from the TOPMed consortium (https://github.com/broadinstitute/gtex-pipeline/blob/master/TOPMed_RNAseq_pipeline.md), with parameters:, with parameters: STAR -- soloBarcodeReadLength 0 --runThreadN 10 --outFileNamePrefix $out. --genomeLoad NoSharedMemory --runRNGseed 789727 --readFilesCommand gunzip -c --outSAMattributes NH HI nM AS CR CY CB UR UY UB sM GX GN --genomeDir $star_index --outSAMtype BAM SortedByCoordinate --outBAMsortingBinsN 200 --outSAMunmapped Within KeepPairs --sjdbGTFfile $gtf --soloType Droplet --soloUMIlen 12 --soloFeatures Transcript3p Gene GeneFull GeneFull_ExonOverIntron GeneFull_Ex50pAS SJ Velocyto --soloMultiMappers Uniform PropUnique EM Rescue --soloUMIfiltering MultiGeneUMI --soloCBmatchWLtype 1MM_multi_pseudocounts --soloCellFilter None --soloCBwhitelist $whitelist–readFilesIn $fastq_files. We filtered the output BAM file using samtools^37,38^ (v. 1.14; options -q 255 -F 4 -F 256 -F 2048). We removed ambient RNA using cellbender^39^ (v. 0.3.0, options --fpr 0.05). Per-nucleus QC metrics were calculated using a custom python script. Code for these steps can be found at https://github.com/porchard/snRNAseq-NextFlow (commit 62e0bdd).

For the chromatin accessibility (ATAC) modality of the multiome data, we trimmed sequencing adapters using cta (v. 0.1.2) and mapped reads to hg38 with bwa^40^ (v.0.7.15; bwa mem -I 200,200,5000 -M). Cell barcodes were corrected using a custom python script implementing the barcode correction algorithm described on the 10x Genomics website. Duplicates were marked using picardtools (http://broadinstitute.github.io/picard/; v 2.26.9; MarkDuplicates READ_ONE_BARCODE_TAG=CB READ_TWO_BARCODE_TAG=CB VALIDATION_STRINGENCY=LENIENT). BAM files were filtered to properly mapped, non-duplicate autosomal read pairs using samtools (samtools view -h -b -f 3 -F 4 -F 8 -F 256 -F 1024 -F 2048 -q 30 chr{1..22}). QC metrics were generated using ataqv^41^ (v. 1.3.0; --ignore-read-groups --nucleus-barcode-tag CB, with the blacklist from ENCODE (https://www.encodeproject.org/files/ENCFF356LFX/)). Code for these steps can be found at https://github.com/porchard/snATACseq-NextFlow (commit 89ab7d5).

### 5’ GEX data processing

The gene expression modality of the single nucleus multiome data was processed using STARsolo^36^ (v. 2.7.10a, with hg38 fasta file and GENCODE v30-derived genome annotations from the TOPMed consortium (https://github.com/broadinstitute/gtex-pipeline/blob/master/TOPMed_RNAseq_pipeline.md), with parameters:, with parameters: STAR --soloBarcodeReadLength 0 --runThreadN 10 --outFileNamePrefix $out. --genomeLoad NoSharedMemory --runRNGseed 789727 --readFilesCommand gunzip -c --outSAMattributes NH HI nM AS CR CY CB UR UY UB sM GX GN --genomeDir $star_index --outSAMtype BAM SortedByCoordinate --outBAMsortingBinsN 200 --outSAMunmapped Within KeepPairs --sjdbGTFfile $gtf --soloType CB_UMI_Simple --soloUMIlen 10 --soloFeatures Transcript3p Gene GeneFull GeneFull_ExonOverIntron GeneFull_Ex50pAS SJ Velocyto --soloMultiMappers Uniform PropUnique EM Rescue --soloUMIfiltering MultiGeneUMI --soloCBmatchWLtype 1MM_multi_pseudocounts --soloCellFilter None --soloCBwhitelist $whitelist --clip5pNbases 39 0 --soloBarcodeMate 1 --readFilesIn $fastq_files. We filtered the output BAM file using samtools^37,38 37,38^ (v. 1.14; options -q 255 -F 4 -F 256 -F 2048). We removed ambient RNA using cellbender^39^ (v. 0.3.0, options --fpr 0.05). Per-nucleus QC metrics were calculated using a custom python script. Code for these steps can be found at https://github.com/porchard/snRNAseq-NextFlow (commit 62e0bdd).

### ONT data processing

The ONT data was processed using a custom NextFlow pipeline (https://github.com/porchard/ONT-snRNAseq-NextFlow, commit 4467492) that borrows from the official ONT wf-single-cell NextFlow pipeline (https://github.com/epi2me-labs/wf-single-cell) but contains several adaptations. Full length reads are identified, trimmed, and oriented, barcodes and UMIs parsed out, and reads mapped in an identical manner as in the official pipeline. The pipelines diverge after the mapping step.

To correct non-whitelisted barcodes, we use the following algorithm. If only one observed whitelisted barcode is within edit distance 2 of the non-whitelisted barcode and all other observed whitelisted barcodes have edit distance greater than 3 from the non-whitelisted barcode, the non-whitelisted barcode is corrected to the whitelisted barcode. Otherwise, we find all observed whitelisted barcodes within edit distance 3 of the non-whitelisted barcode and calculate the relative probability that the non-whitelisted barcode derived from each (first, we calculate the probability of the observed errors between each and the non-whitelisted barcode using the barcode phred scores, or error probability = 0.05 in the case of an indel; next, we weight each of these values by the number of times each of the whitelisted barcodes was observed in the data; last, we normalize the resulting values to sum to one). We correct the non-whitelisted barcode to the most probable whitelisted barcode if its relative probability exceeds 0.95.

We assign reads to genes based on gene body overlap (requiring same strand). If a read overlaps the gene body of a single gene, it is assigned to that gene. If it overlaps the gene body of more than one gene it is assigned to the gene with exon overlap, or left unassigned (it if doesn’t overlap any exons or overlaps exons of more than one gene. We assign reads to transcripts using IsoQuant^42^ v. 3.5.2 (--output. --bam $bam --data_type ont --reference $fasta -- no_model_construction --complete_genedb --genedb $gtfdb --stranded none --no_secondary -- labels $library --bam_tags CR,UR --prefix $library --genedb_output genedb/; $gtfdb was generated using gffutils v. 0.13 with command gffutils.create_db($gtf, $gtfdb, disable_infer_genes=True, disable_infer_transcripts=True)); IsoQuant output is filtered to keep cases where assignment type is one of ‘unique’, ‘unique_minor_difference’, ‘inconsistent’, or ‘inconsistent_non_intronic’.

UMIs are corrected using the same algorithm as in the original ONT pipeline, using the barcode corrections and gene assignments from our custom pipeline.

To remove ambient RNA we run CellBender^39^ v. 0.3.0 (cellbender remove-background --cuda -- epochs 150 --fpr 0.05).

### Sample genotyping

We conducted light whole-genome sequencing (WGS) on DNA isolated from the muscle biopsies. SNPs were imputed with GLIMPSE2 using 1000 Genomes as a reference and filtered for HWE < 1e-6 and MAF >= 5%.

### Nuclei QC

After multiome snRNA and snATAC preprocessing, we applied several QC thresholds on both RNA and ATAC barcodes. For RNA barcodes, we considered (using the Illumina-sequenced 3’ GEX): minimum UMI = 120, maximum mitochondrial read fraction = 0.1, minimum probability of singlet from cellbender = 0.99, post-cellbender UMIs = 100, maximum fraction of exon vs full-gene-body reads = 0.6. For ATAC barcodes, we considered library-specific minimum high-quality autosomal alignments (HQAA) thresholds, which ranged from 3000 to 10000, and minimum TSS enrichment = 2.

For 5’ GEX, we used the following thresholds: minimum UMI = 120, maximum mitochondrial read fraction = 0.1, minimum probability of singlet from cellbender = 0.99, post-cellbender UMIs = 100, maximum fraction of exon vs full-gene-body reads = 0.6

### Genetic demultiplexing

Prior to running demuxlet, 3’ GEX BAM files were filtered to uniquely mapped alignments (samtools view -h -b -q 255 -F 4 -F 256 -F 2048 $bam).

We ran demuxlet (popscle commit da70fc7) on pass-QC barcodes with the following parameters:

popscle demuxlet --sam $bam --tag-group CB --tag-UMI UB --min-mac 1 --vcf $vcf --sm-list $sample_list --out $out --group-list $barcode_list

For RNA, the input VCF file was filtered to variants overlapping gene bodies (based on the GENCODE v30 annotation).

For ATAC, the input VCF file was filtered to variants in TSS-distal ATAC peak summits from a previous single-nucleus muscle dataset^30^.

When comparing demuxlet assignments across modalities and sequencing technologies, we recoded ambiguous (‘AMB’) droplet types as doublets (‘DBL’).

### SCAFE

Prior to running SCAFE, 5’ GEX BAM files were filtered to uniquely mapping (mapping quality 255) primary alignments (samtools view -h -b -f 1 -F 2316 -q 255). They were then preprocessed for compatibility with SCAFE as described at https://github.com/chung-lab/SCAFE/issues/29.

ONT BAM files were filtered to mapped primary alignments with minimum mapping quality 10 (samtools view -h -b -F 2308 -q 10). We also preprocess ONT BAM files with custom python scripts prior to running SCAFE, as described below (for “ONT GEX (trim only)” processing the below steps are skipped, except for read trimming from the 3’ end which is required to avoid a SCAFE error).

Alignments from the official ONT single cell pipeline, as well as our custom pipeline, are expected to begin with a guanine trinucleotide sequenced (GGG), representing the last 3 basepairs of the TSO sequence. Transcripts with 5’ mRNA caps may have a fourth guanine, representing the cap. Therefore, we filter the BAM files as follows. First, the amount of softclipping at the beginning of the read is determined, and the read is skipped if there are more than 4 bps of softclipping (which allows for a possibly unaligned ‘GGG’ + ‘G’ sequence). Next, the 5’ end of the read is checked for the residual ‘GGG’ from the TSO, allowing for one mismatch; if it is not present, the read is filtered out, and if it is present, the three bps at the 5’ end of the read are trimmed. Then, the read is trimmed from the 3’ end until it is no longer than 100 bps.

In addition, all PCR duplicates should nominate the same 5’ transcript end. However, artifacts in library preparation or sequence alignment may introduce discrepancies between apparent duplicate reads. Therefore, before running SCAFE, we identify groups of reads with the same cell barcode (CB tag), UMI (UB tag), and gene assignment (GN/GX tag), and check the 5’ end positions. If they do not all nominate the same 5’ end position, we determine the 5’ end position with the most reads supporting it and keep only the corresponding reads, filtering out the others. If two 5’ end positions have equal support, we keep the most upstream one.

We use SCAFE^32^ v1.0.0, a suite of PERL-based scripts, to quantify reads associated with 5’ ends of sequenced molecules. First, BAM files are processed to extract 5’ end counts (cTSS) using the tool.sc.bam_to_ctss command. Importantly, we define the template switch oligo specific to 10x 5’ vs. 3’ kits (--TS_oligo_seq) and set --detect_TS_oligo ‘auto’, except for the ONT GEX (trim only) case for which --TS_oligo_seq is set to ‘GGG’ and --detect_TS_oligo is set to ‘match’ (reflecting the fact that the rest of the TSO sequence is removed in upstream processing). Next, strand invaders are identified and removed from cTSS counts using the command tool.cm.remove_strand_invader with a minimum edit distance of 5 (--min_edit_distance=5) and a minimum number of upstream non-G nucleotides of 2 (--min_end_non_G_num=2); --TS_oligo_seq for this step is set according to the appropriate TSO sequence from the 3’ or 5’ kit (including for ONT GEX (trim only)). cTSS counts are then clustered using *Paraclu*^43^ with the tool.cm.cluster command including clusters with at least 5 counts across the cluster (--min_cluster_count=5) and at least 3 counts at the cluster peak, (--min_summit_count=3). We select a third parameter, --min_num_sample_expr_cluster (in our case representing the minimum number of nuclei required to support the cluster) separately for each dataset (Table S1) based on examining the number of tCREs discovered in TSS and enhancer chromatin states versus the number discovered in other chromatin states as the parameter is varied, selecting the parameter value at which the rate of discovery of non-TSS, non-enhancer tCREs begins to rapidly increase without a corresponding rapid increase in TSS/enhancer tCREs. Logistic regression is then used to filter clusters based on the number of UMI within 75 nt of the cluster peak, unencoded G percentage, and corrected expression using the command tool.cm.filter with a cutoff of 0.5 (--default_cutoff=0.5). Corresponding ATAC data is used for training the model. Filtered clusters are then annotated using the tool.cm.annotate command to generate a final set of tCREs. Additionally, bigwig files are generated from cTSS counts before and after strand invader removal using the command tool.cm.ctss_to_bigwig.

### GAT

tCRE enrichment in chromatin states were calculated using GAT^44^ (v. 1.3.6), with command gat-run.py --ignore-segment-tracks --segments=$tcres --annotations=chromatin-states.bed -- workspace=hg38.workspace --num-samples=10000, where hg38.workspace consists of autosomes.

### Chromatin states

Skeletal muscle chromatin states were from Varshney et al.^33^, lifted from hg19 to hg38 using liftOver^45^.

