## Supplementary Figures 1-5 for "Genetic demultiplexing and transcript start site identification from nanopore sequencing of 10x Genomics multiome libraries"

**A.**

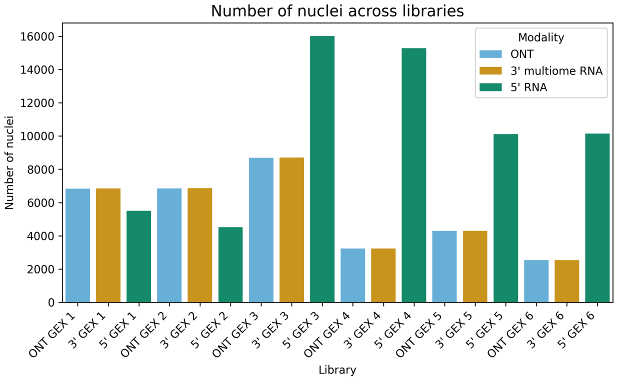


**B.**

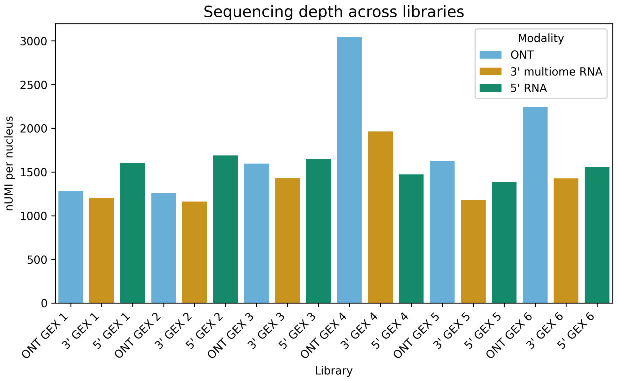


**C.**

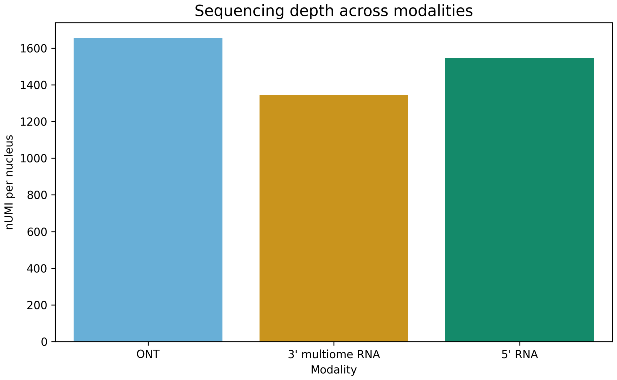


**Fig. S1.** (A) Number of nuclei per library. (B) Mean number of UMIs per nucleus per library. (C) Mean number of UMIs per nucleus per modality.

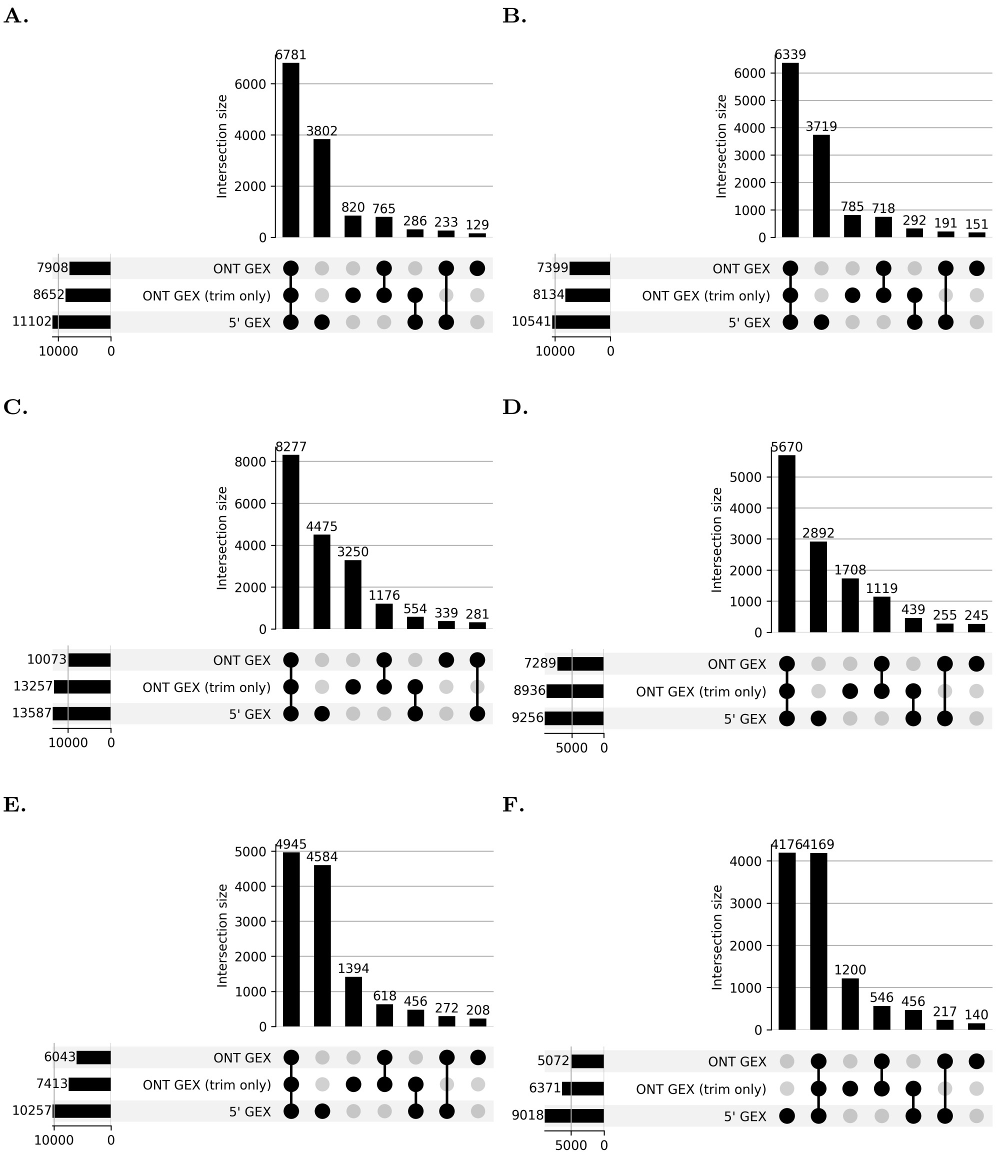


**Fig. S2.** Upset plots showing tCRE overlaps between 5' GEX, ONT GEX (trim only), and ONT GEX. ONT GEX includes the full processing outlined in the left panel of Fig. 2A, whereas ONT GEX (trim only) includes only trimming of reads from the 3' end prior to running SCAFE.

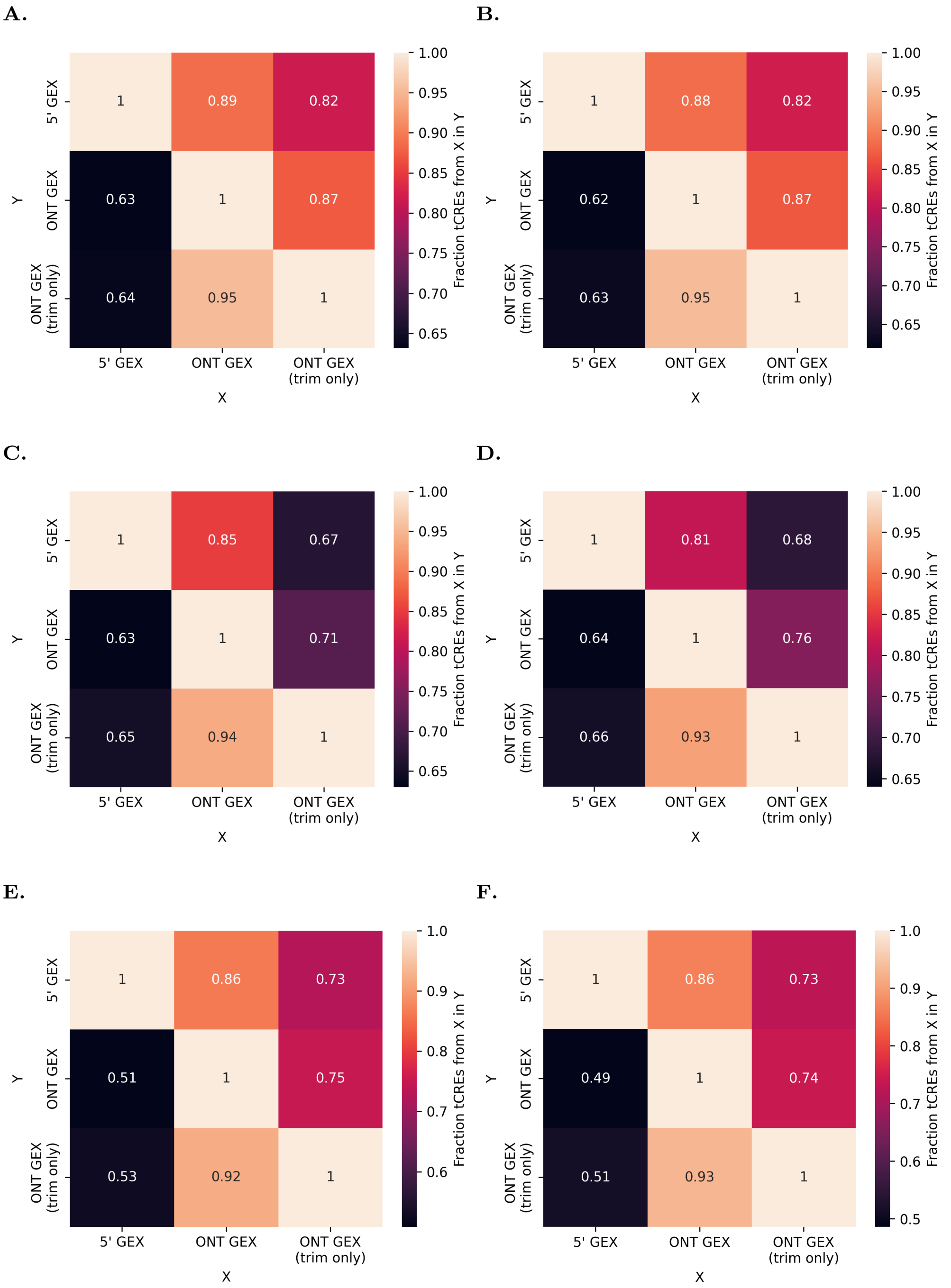


**Fig. S3.** Pairwise comparisons of tCRE overlaps between 5' GEX, ONT GEX (trim only), and ONT GEX. ONT GEX includes the full processing outlined in the left panel of Fig. 2A, whereas ONT GEX (trim only) includes only trimming of reads from the 3' end prior to running SCAFE.

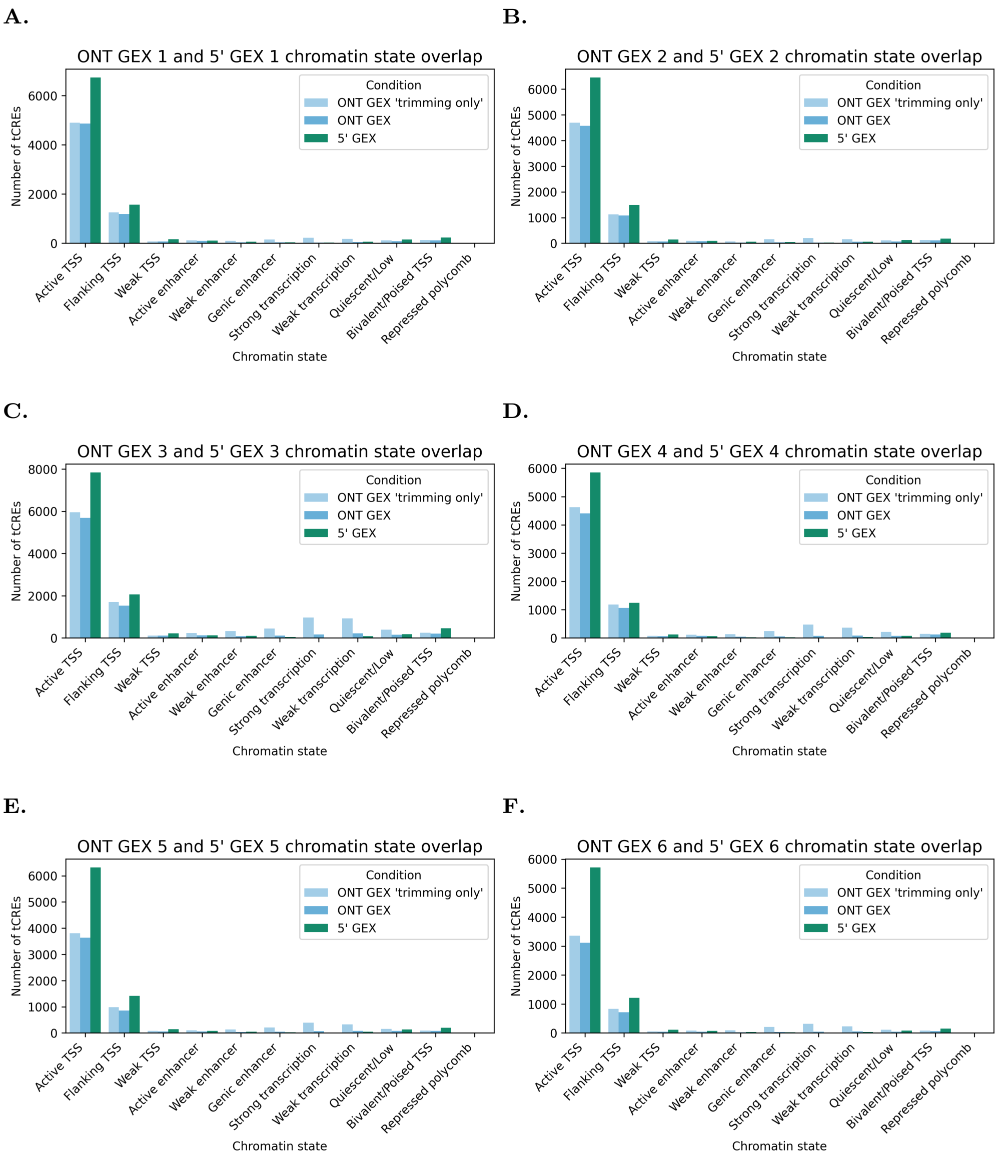


**Fig. S4.** tCRE chromatin state overlaps for 5' GEX, ONT GEX (trimming only), and ONT GEX. ONT GEX includes the full processing outlined in the left panel of Fig. 2A, whereas ONT GEX (trimming only) includes only trimming of reads from the 3' end prior to running SCAFE.

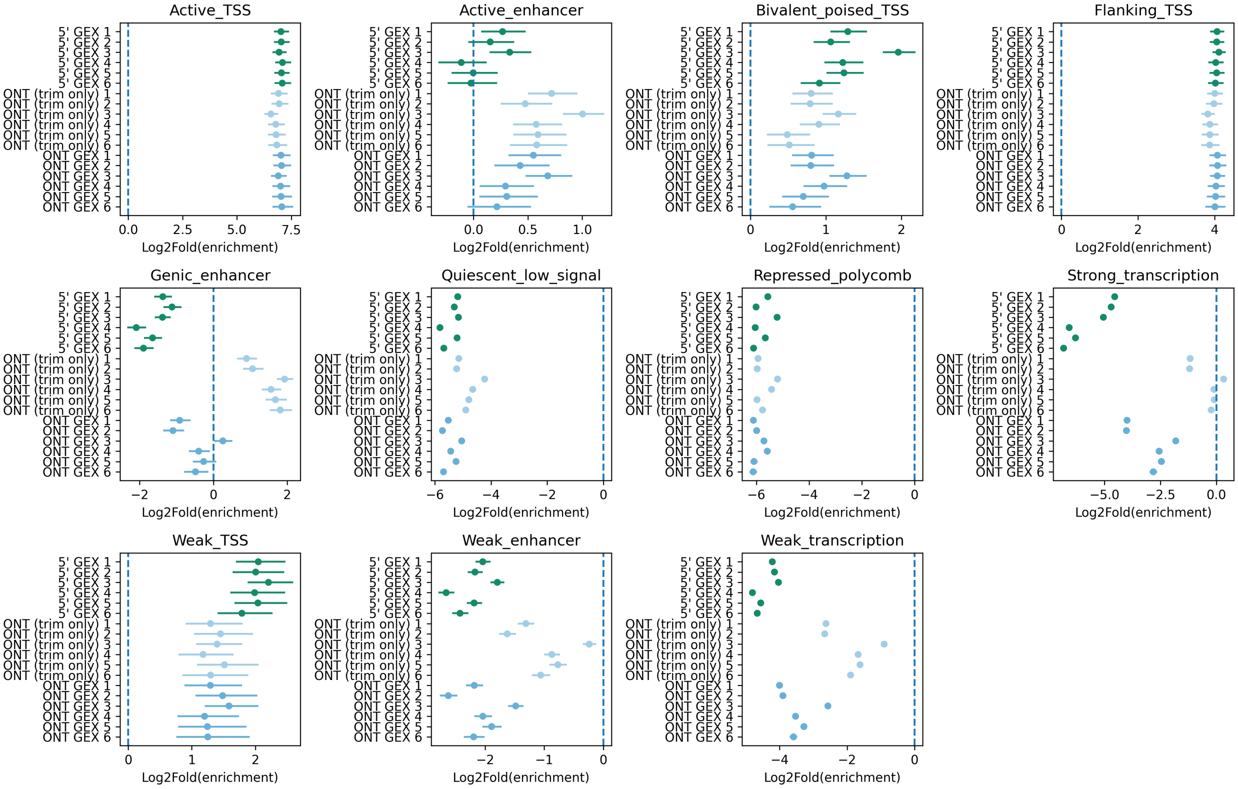


**Fig. S5.** tCRE enrichment in chromatin states.
