## Supplemental Table 1 for "Genetic demultiplexing and transcript start site identification from nanopore sequencing of 10x Genomics multiome libraries"

| Library | Cluster counts | Summit counts | Number of nuclei |
| --- | --- | --- | --- |
| ONT GEX 1 | 5 | 3 | 7 |
| ONT GEX 2 | 5 | 3 | 7 |
| ONT GEX 3 | 5 | 3 | 8 |
| ONT GEX 4 | 5 | 3 | 8 |
| ONT GEX 5 | 5 | 3 | 8 |
| ONT GEX 6 | 5 | 3 | 8 |
| ONT GEX 1 'trim only' | 5 | 3 | 9 |
| ONT GEX 2 'trim only' | 5 | 3 | 9 |
| ONT GEX 3 'trim only' | 5 | 3 | 10 |
| ONT GEX 4 'trim only' | 5 | 3 | 13 |
| ONT GEX 5 'trim only' | 5 | 3 | 13 |
| ONT GEX 6 'trim only' | 5 | 3 | 13 |
| 5' GEX 1 | 5 | 3 | 13 |
| 5' GEX 2 | 5 | 3 | 12 |
| 5' GEX 3 | 5 | 3 | 15 |
| 5' GEX 4 | 5 | 3 | 14 |
| 5' GEX 5 | 5 | 3 | 11 |
| 5' GEX 6 | 5 | 3 | 10 |
